# Linked optical and gene expression profiling of single cells at high throughput

**DOI:** 10.1101/766683

**Authors:** Jesse Q. Zhang, Christian A. Siltanen, Leqian Liu, Kai-Chun Chang, Zev J. Gartner, Adam R. Abate

## Abstract

Single cell RNA sequencing has emerged as a powerful tool for characterizing cells, but not all phenotypes of interest can be observed through gene expression alone. Linking sequencing with optical analysis has provided insight into the molecular basis behind cellular function, but current approaches have limited throughput. Here, we present a high throughput platform for linked optical and gene expression profiling of single cells. We demonstrate accurate fluorescence and gene expression measurements from thousands of cells in a single experiment and use the platform to characterize DNA and RNA changes in Jurkat cells through the cell cycle. In addition to its scalability, our integration of microfluidics and array-based molecular biology holds promise for comprehensive multi-omics profiling of single cells.

## Introduction

Cellular processes such as replication, migration, and differentiation, are tightly controlled by signaling and gene regulatory networks (1,2,3). These processes are dynamic, and at any point a cell may exist along a continuum of states (4). Thus, cell state heterogeneity is often masked when bulk methods are used to analyze populations (5,6). The development of high throughput single cell RNA sequencing (scRNA-seq) has enabled populations to be analyzed at the single cell level (7,8,9), leading to the dissection of cellular heterogeneity and the construction of a map of cell states across the human body (10). However, gene expression is just one dimension by which cells may be characterized, and many properties, such as epigenetic state, protein expression, enzyme activity, and cellular morphology, are not readily measured by scRNA-seq alone (11,12).

More comprehensive cell characterization can be accomplished by combining scRNA-seq with complimentary measurement methods. Optical approaches, including microscopy and flow cytometry, can characterize morphological and fluorescence phenotypes prior to scRNA-seq (13,14). Linked optical analysis and scRNA-seq has been applied to *in vitro* cultures, patient tissues, and stem cells, revealing molecular links to cellular function (15,16,17). While powerful, these approaches are limited in throughput. Imaging methods require cells be imaged and individually transferred to wells for sequencing (15). Cytometry methods are more scalable, since the instrument can automatically sort cells into wells for automated library preparation, increasing throughput to hundreds of cells (18). Scaling beyond this limit, however, is impractical because the time and volume of reagent required to process tens or hundreds of thousands of cells is prohibitive (19). Recent spatial transcriptome sequencing approaches might ultimately enable scalable imaging and scRNA-seq, but they rely on non-standard methods to image and label cells (20,21). To enable scalable optical and scRNA-seq analysis of large populations, a new approach is needed that can rapidly perform optical measurements, then isolate and sequence large numbers of single cells.

In this paper, we present a high throughput platform for linked analysis of optical phenotype and gene expression of single cells. Our instrument functions like a flow cytometer, optically scanning cells in flow and dispensing them to a well plate where they are prepared for sequencing. The optical analysis is accomplished using a microfluidic-based droplet cytometer, and the cells are dispensed into custom nanoliter volume wells. Once isolated, cellular mRNAs are indexed and prepared for sequencing such that each sequenced read contains information about the well from which it originated; this allows all reads to be assigned to their original wells, thereby providing a linked dataset of optical and RNA-seq information for each single cell. The cell analysis is accomplished at ∼1 KHz and dispensing to the array at ∼5 Hz, allowing a thousand cells to be isolated in a few minutes. The total volume of reagent on the array is ∼1 µL per 1,000 single cell transcriptomes, representing a 1000-fold reduction in reagent volume compared to microwell plates, affording a significant cost savings. Moreover, with simple fabrication techniques, ∼10,000 wells can be fabricated on a chip with a footprint comparable to a standard microscope slide. Thus, our approach provides a scalable means by which to acquire linked single cell optical and gene expression data for large cell populations.

## Results and Discussion

Our single cell analysis platform is based on Printed Droplet Microfluidics (PDM) (22,23), an approach that allows cells to be optically scanned and dispensed to custom nanoliter volume well plates (nanoplates) (**Fig. 1A**). To perform linked optical and scRNA-seq analysis, we record the fluorescence of a cell while confined to a droplet, then dispense the cell and droplet to the nanoplate at defined locations. Then, scRNA-seq library preparation is performed on each cell using specific “coordinate oligos” encoding each cell’s location on the nanoplate. After sequencing, these oligos allow each cell barcode to be traced to a well of origin, thereby linking it to the optical data collected for that cell. The workflow is similar to flow cytometry, except that the sorter is a microfluidic device and the wells in which the cells are dispensed are ∼10,000-fold smaller than conventional microwells (∼100 pL). This reduction in volume, combined with the speed of the microfluidic printer, enable highly scalable optical phenotyping and sequencing of single cells.

**Figure 1.**
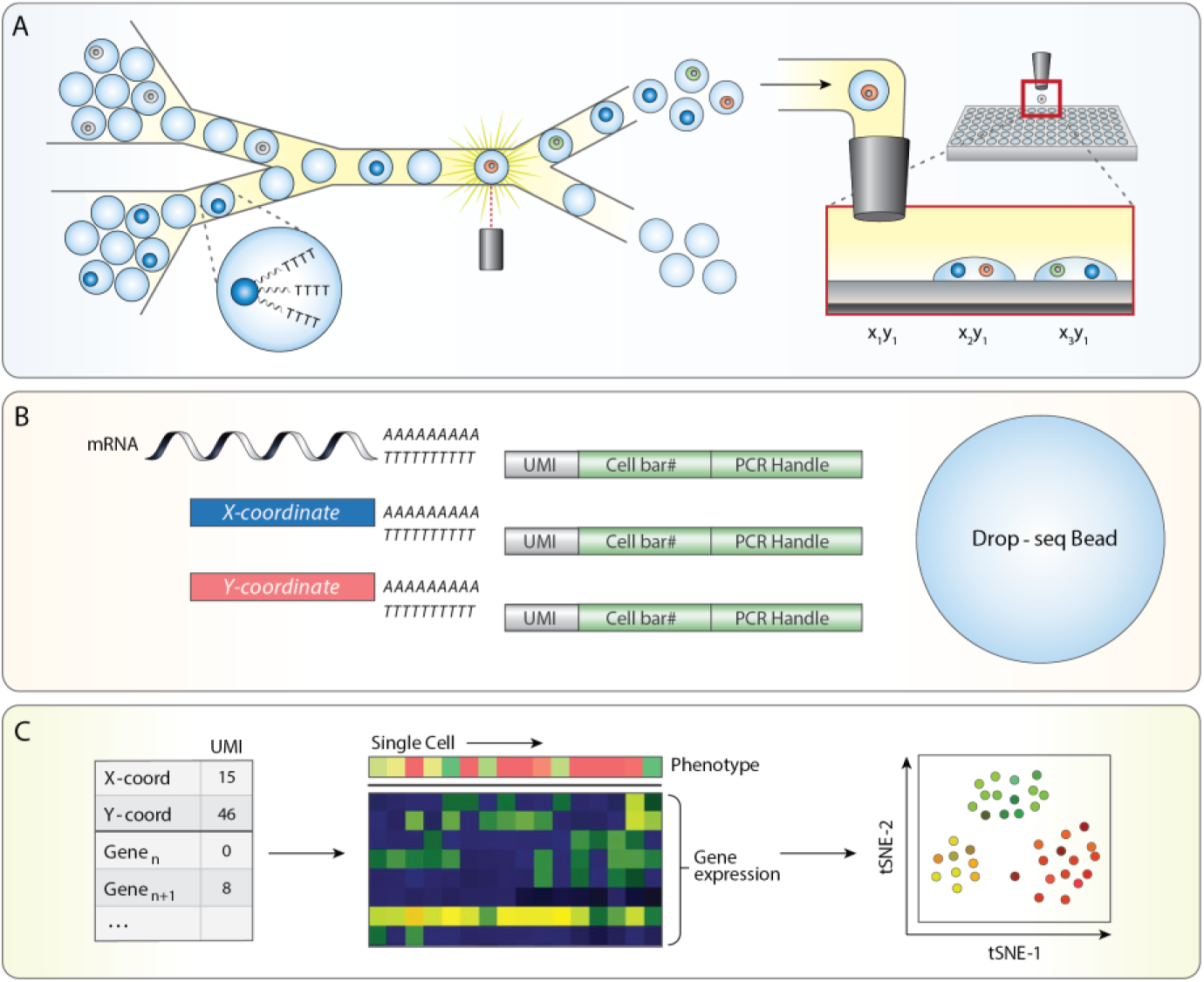
A high throughput platform for linked optical phenotype and gene expression of single cells. (A) Monodisperse droplet emulsions containing encapsulated poly-T mRNA capture beads and cells are input into a microfluidic device. Fluorescence signal from droplets is interrogated and used to selectively dispense a cell and bead to indexed locations on a nanowell array. (B) Each bead binds mRNA from cell lysate as well as a unique combination of poly-A barcode oligos denoted by nanowell coordinate. (C) UMI counts on each bead are collected through sequencing into an expression matrix for each cell. Nanowell coordinate is assigned based on the abundance of barcode oligos and paired with fluorescence data obtained during cell sorting, which enables downstream linked analyses such as dimensionality reduction visualizations of gene expression paired with optical phenotype.

Prior to device operation, a separate flow focusing device encapsulates cells in a droplet emulsion. We introduce this emulsion into the PDM device, where each drop is optically scanned (**Fig. 1A, left**). As in flow cytometry, laser-induced fluorescence accomplishes the optical analysis, whereby focused lasers excite fluorescence of the cells which a multicolor detector then captures (**Fig. 1A, middle**). Cells with desired fluorescence properties are isolated through sorting them into a printing nozzle that dispenses them into a nanowell on the substrate (**Fig. 1A, right**). We record the cell fluorescence and dispense location, allowing this information to be paired with the scRNA-seq data collected later.

To link the optical and sequencing data, we index the wells such that each cells’ dataset can be traced back to a well on the array. The indexes comprise “coordinate oligos” pre-loaded into the wells using a commercial reagent spotter (**Fig. 1B, lower left**) (24). To index the array, we place “X” and “Y” coordinate oligos, each of which contains a different 8 base sequence encoding the specific location of a given nanowell on the plate. The coordinate oligos are polyadenylated, allowing them to be captured with cellular mRNA during the scRNA-seq library preparation. For scRNA-seq, we adapt the validated “Drop-Seq” protocol (7), which uses beads coated with poly-thymine “barcode” oligos to capture and label both mRNA and coordinate oligos. We accomplish this by co-dispensing beads and cells in each nanowell and lysing the cells. After retrieving the beads, performing the requisite library preparation steps of Drop-Seq, and sequencing the barcoded cDNA, we obtain a collection of reads representing the cell transcriptome and coordinate oligos, all sharing a Drop-Seq barcode. Thus, the location of the cell from which the data originate is encoded in the sequencing data, allowing it to be traced back to a well on the array (**Fig. 1C, left**) and associated with the previously recorded optical data (**Fig. 1C, middle**). With this paired dataset, we can use dimensionality reduction methods to first visualize gene expression data, to which we add the optical phenotype information (**Fig. 1C, right**).

The microfluidic print device consists of a droplet spacer, sorter, and printing nozzle (**Fig. 2A**). A packed emulsion containing cells or beads is introduced, spaced by oil, and optically scanned by a four-color laser-induced fluorescence detector (**Fig. 2A, red outline**). Embedded fiber optics excite and collect fluorescence that is processed through filters and analyzed in real time by custom software; this allows cell, bead, and droplet fluorescence and scattering data to be recorded, to determine whether to print the droplet and its contents to the current nanowell. Printing is achieved by sorting a droplet (**Fig. 2A, green outline**) into the printing nozzle positioned above a nanowell (**Fig. 2A, purple outline**); if the current droplet should not be printed, it is not sorted into the nozzle and instead passes into the discard channel. Because the carrier oil is viscous and denser than water, in the absence of other forces, the ejected droplet would float away and not go into the nanowell. Thus, to dispense it into the nanowell, electrodes positioned under the substrate emit an oscillating electric field. This field pulls the dispensed droplet into the nanowell and is key to the speed of PDM, since it allows a droplet to travel the final few hundred microns from the printing nozzle to the nanowell in tens of milliseconds (22). Moreover, because the trap extends above the substrate, the printer need not dispense the droplets with perfect accuracy into the nanowells, since any droplet within the electric field will, ultimately, be pulled into the nearest nanowell. The trapping field also ensures that the printed droplets remain fixed in the wells. Upon completion of a print run, droplets can be released by un-powering the electrodes (**Movie S1**).

**Figure 2.**
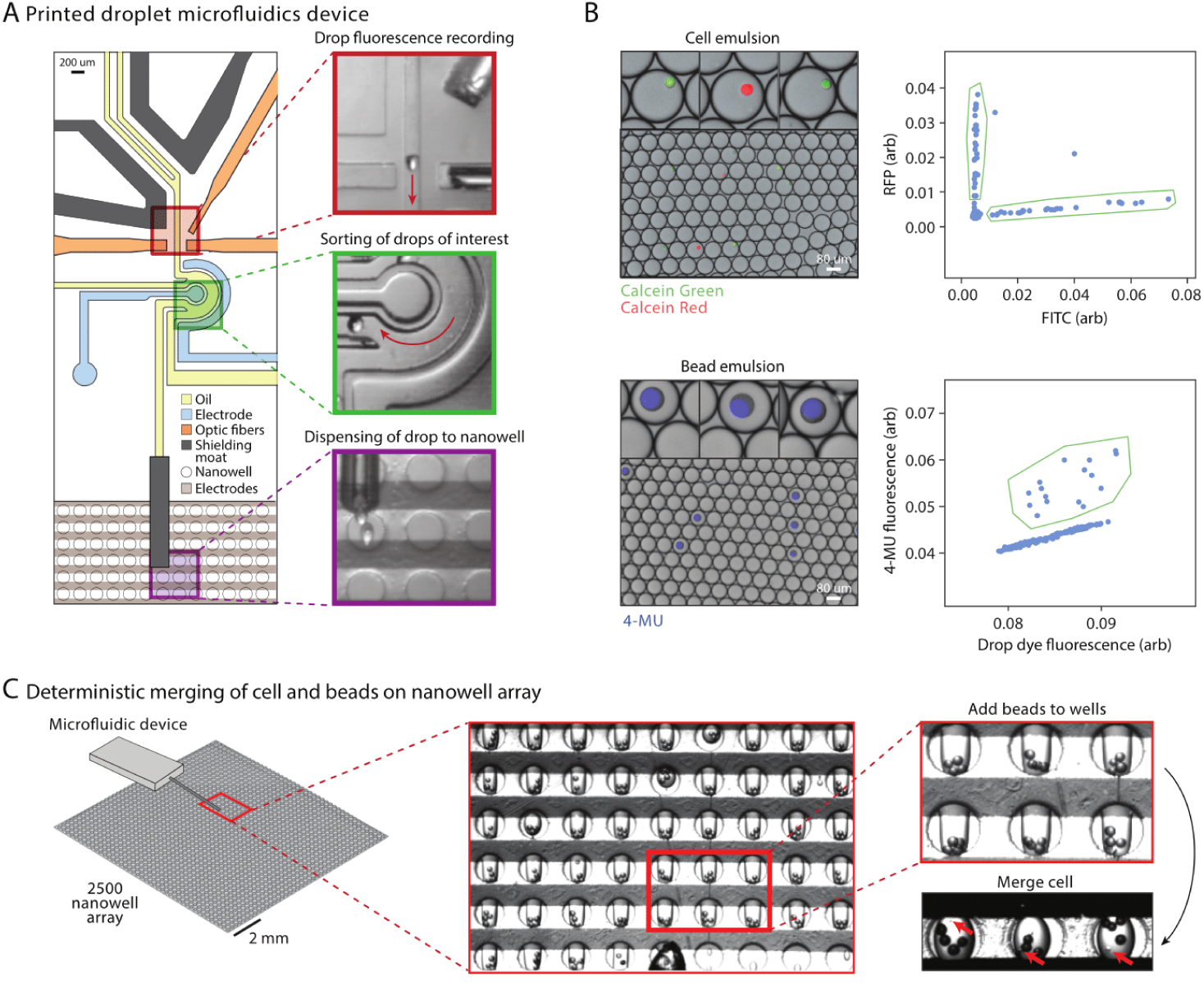
Printed Droplet Microfluidics (PDM) operation for deterministic loading of nanowell array with beads and cells. (A) An inset of the microfluidic device aligned over the nanowell array, with images (top to bottom) of regions of drop fluorescence recording, sorting of drops of interest, and dispensing of drops to nanowells. (B) Monodisperse droplet emulsions containing fluorescently labeled cells (top) or beads (bottom) are input into PDM. Drops of interest (insets) are enriched for by gating on fluorescence plots (right) generated during device operation. (C) Deterministic merging of cells and beads through first adding beads to nanowells, followed by merging of a cell-containing drop in lysis buffer.

To demonstrate the accuracy of scRNA-seq using our approach, we perform a two-cell experiment. We prepare and encapsulate a mixed suspension of Calcein Red stained mouse (3T3) and Calcein Green stained human (HEK293) cells (**Fig. 2B, upper left**). When scanned in the print head, we observe distinct green and red cell populations (**Fig. 2B, upper right**); thus, with suitable gating instructions, the printer can print these cells in a defined pattern to the nanoplate. To enable scRNA-seq of the printed cells, Drop-Seq beads must also be printed, which requires that they be detectable in the print head; this is accomplished by labeling them with 4-MU, a blue dye that does not overlap with the cell stains (**Fig. 2B, lower left**). This signal allows bead-loaded droplets to be discerned from bead-empty droplets (**Fig. 2B, lower right**). To print cells and beads in defined combinations, we generate a “print file” containing gating and location instructions that we input into the printer software; the printer reads this file, printing cells and beads to the nanoplate according to the instructions in the file (**Fig. 2C, left)**.

A strength of PDM for scRNA-seq is that it allows the nanowells to be controllably loaded with cells and beads; this contrasts with other high throughput scRNA-seq methods which randomly load cells or beads and, thus, are less efficient. Moreover, PDM allows systematic variation of nanowell contents across the array, to choose conditions that maximize data quality (22). For example, controlled cell loading minimizes doublets, nanowells in which two cells are inadvertently sequenced as one, which can be major confounders in scRNA-seq experiments that assume single-cell data (25,26). Moreover, controlled printing allows us to load multiple capture beads to every well (**Fig. 2C, right, Movie S2**), which can increase cell and mRNA capture efficiency by compensating for losses during sequencing library preparation (27). To illustrate this, we print two substrates, the first with one bead per well and the second with four, both on 42 by 56 nanowell (2352) arrays. All beads originating from the same well contain the same coordinate barcodes, allowing us to group reads associated with multiple beads together. Due to loss of beads during library preparation, starting with more beads per well increases the likelihood of recovering at least one bead from every well (**Fig. 3A**). When printing more beads, we also recover more transcripts per well (**Fig. 3B, upper**), and that the number of transcripts per bead remains consistent when recovering up to 5 beads per well (**Fig. 3B, lower**). This suggests that increasing bead surface area per cell lysate increases mRNA capture.

**Figure 3.**
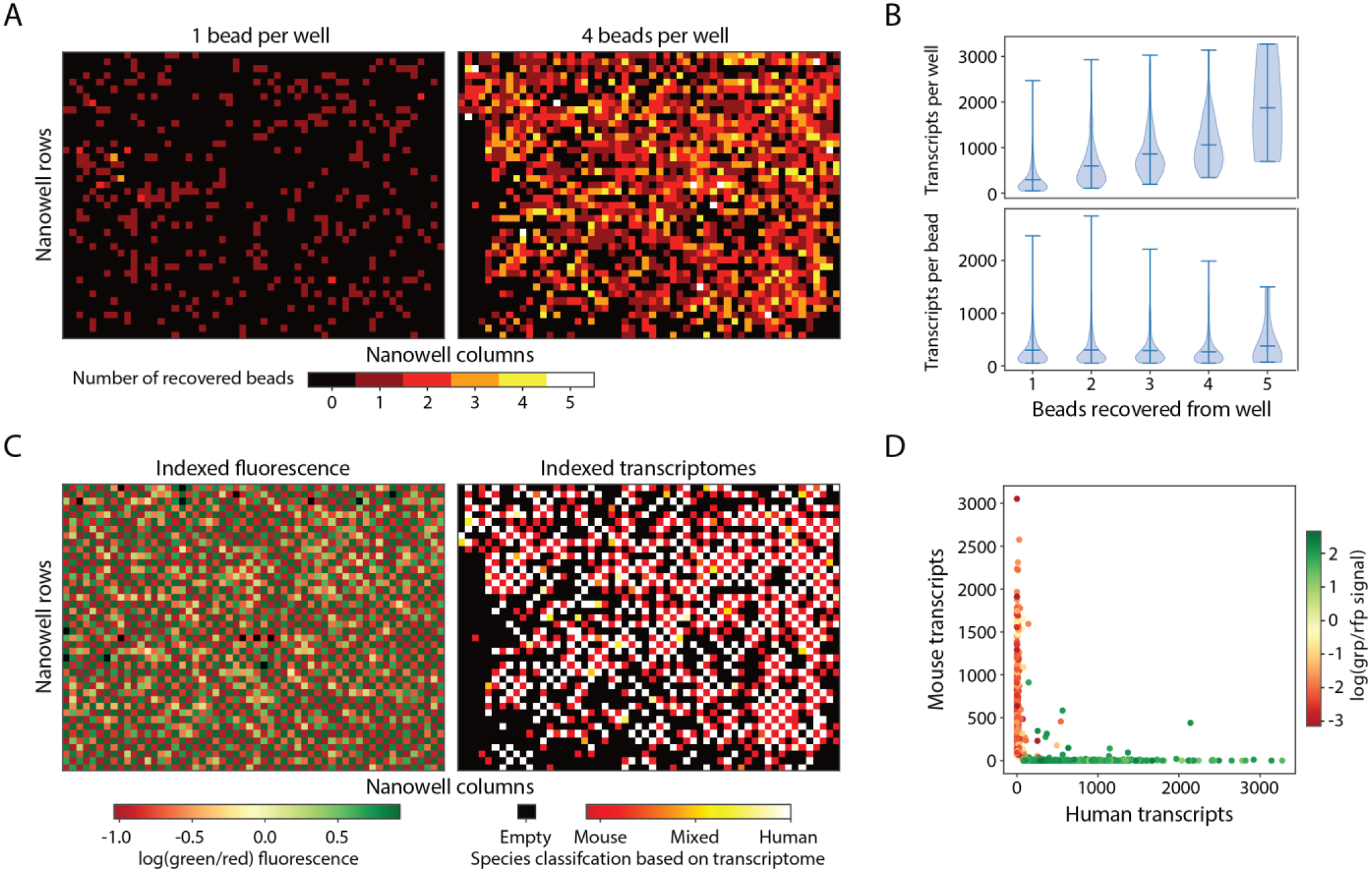
Linked optical phenotype and gene expression measurements verified with two species experiment. (A) One or four beads were printed to each well of a 42 by 56 nanowell array along with an alternating pattern of mouse and human cells. The number of recovered beads per nanowell position was determined by the number of unique cell barcodes mapped back to each nanowell. (B) When printing four beads per well, the distribution of transcripts recovered from each nanowell were calculated as a function of the number of beads recovered. The distribution of the number of transcripts originating from each bead within a nanowell was also plotted as a function of the number of beads recovered per nanowell. (C) Left: Fluorescence data from alternating printing of Calcein green stained human cells and Calcein red stained mouse cells indexed by nanowell position. Right: Ratio of human to mouse transcripts recovered from each nanowell based on printing four beads per nanowell. (D) Transcript counts by nanowell position are annotated with the green-red fluorescence ratio from the cell printed into the corresponding nanowell.

Single cell RNA-seq methods require minimal cross contamination of RNA between cells (25). To investigate cross contamination, we print mouse and human cells in a checkerboard pattern and classify each transcriptome according to which species’ genome the transcripts predominately align. We find that the optical data align 99.3% of the time with the expected printing pattern (**Fig. 3C, left**). We recover transcripts from 1191 nanowells, with 94.3% of transcriptomes having species purities of at least 90% mouse or 90% human and matching the expected printing pattern (**Fig. 3C, right**). We find that 5.3% of nanowells have less than 90% species purity, suggesting either mRNA cross contamination or misprinting of cells (dots not aligned with axes) (**Fig. 3D**). In total, we recover quality optical and scRNA-seq data from over a thousand cells when printing four beads per well.

Cells undergo changes in state and phenotype through the cell cycle (17). For example, by the G2 phase, cells have doubled their genome, allowing it to be optically detected via DNA staining (28). Moreover, different genes peak and diminish in expression through the cycle, making the cell cycle useful for validating our approach. As a model system, we use Jurkat cells stained with DRAQ5, which is a live-cell stain for genomic DNA (**Fig. 4A, upper**). We observe a broad distribution of DNA fluorescence, the brightest of which likely correspond to cells with the most DNA and, thus, in G2M-phase. To confirm these results, we perform flow-cytometry of the suspension and obtain a similar distribution (**Fig. 4A, lower**). We utilize PDM to generate a 56 by 56 nanowell array to which we print a checkerboard pattern of low and high DRAQ5 expressing cells (**Fig. 4B**). We sequence transcriptomes from 437 cells and use Uniform Manifold Approximation and Projection (UMAP) to visualize these cells (29). Through assigning cells to G1, S, or G2M phases based on their expression of cell cycle-associated genes and using those genes for principal component analysis, we generate a UMAP plot which identifies three clusters in agreement with three stages of the cell cycle (**Fig. 4C, left**). To determine whether these classifications agree with the optical data, we annotate the points of the UMAP plot according to the magnitude of DRAQ5 fluorescence (**Fig. 4C, right**). The plots are in general agreement, with the state comprising the most DNA (G2M) appearing brightest in the DNA stain. To observe how the population varies through this cycle, we order the cells by fluorescence and plot the proportion in the three states as classified by gene expression (**Fig. 4D**). We expect the proportion of cells in G1 to be at low DRAQ5 signal, S phase in the middle of the distribution, and G2 at the top end. The peak of the G1 phase curve is at the low end of the DRAQ5 distribution, S-phase in the middle, and G2/M at the top end. We observe general concordance with the expected trend when we pair fluorescence and scRNA-seq measurements of cell cycle state. With our platform we thus demonstrate characterization of a fundamental biological process through linked optical and gene expression analysis.

**Figure 4.**
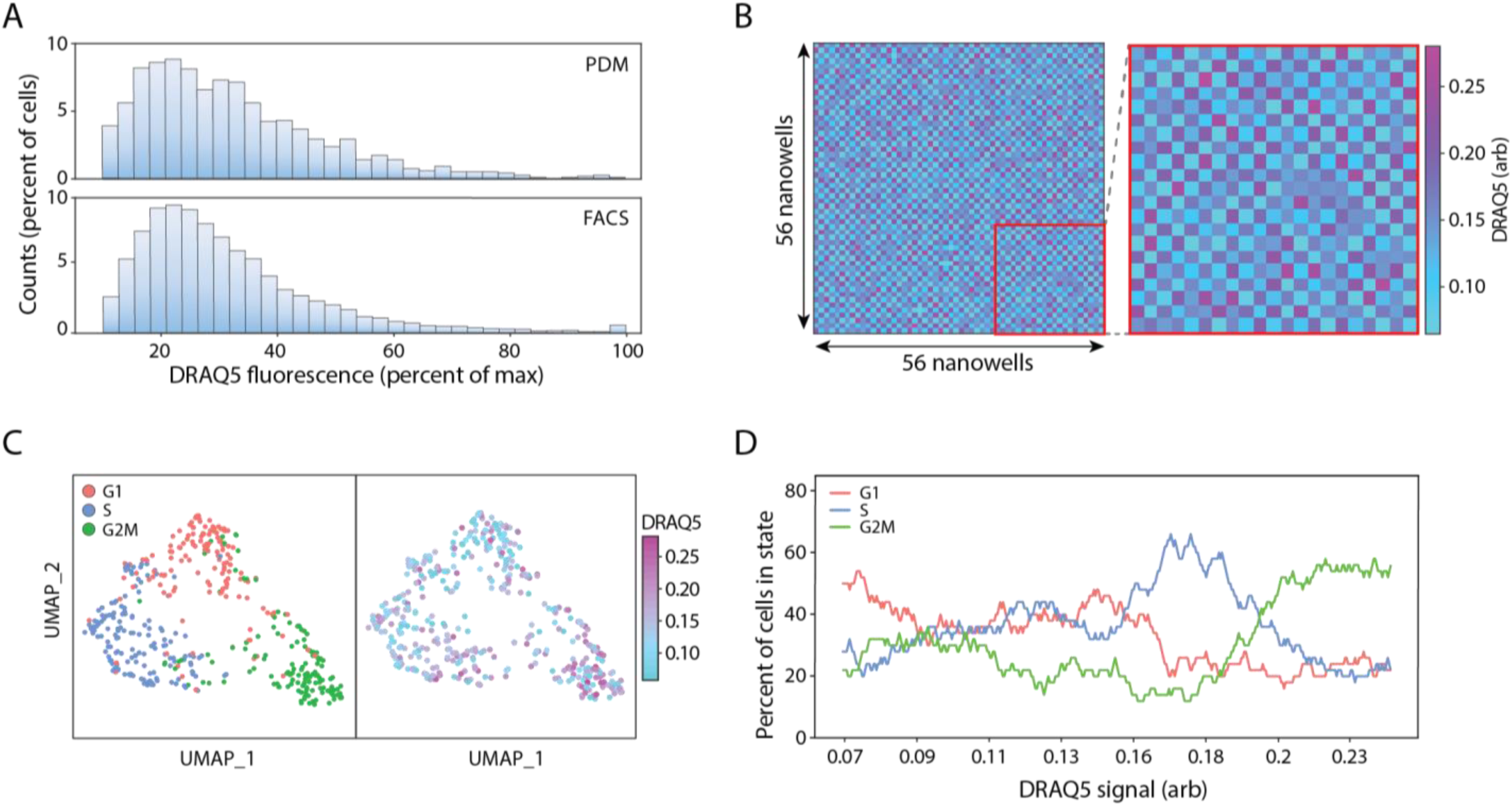
Linked fluorescence and gene expression analysis of cell cycle state in Jurkat cells stained with a DNA-binding dye. (A) The frequency distribution of Jurkat cells stained with DRAQ5 encapsulated within droplets was analyzed on both PDM and a flow cytometer. (B) An alternating pattern of high and low expressing DRAQ5 Jurkats was dispensed to a 56 by 56 nanowell array using PDM. Fluorescence measurements were indexed by nanowell position (inset). (C) Transcriptomes from 498 cells were recovered and clustered based on cell cycle state (left) predicted based on a set of cell cycle dependent genes. DRAQ5 fluorescence data collected during printing was then overlaid (right). (D) Cells were ordered by low to high DRAQ5 signal (top bar), and the fraction of cells in each cell cycle state was calculated over a 50-cell sliding window using corresponding cell cycle state assignments by gene expression analysis.

Single cell RNA-seq is a powerful and general method for analyzing cells, but not all traits of interest are observable in gene expression data alone. Here, we demonstrate an approach that allows gene expression to be linked to optical data in a high-throughput format scalable to thousands of single cells. A key concern for any scRNA-seq workflow that relies on barcoding cells for bulk sample amplification, is the loss of cells due to loss of beads during downstream processing (27). By spreading each cell’s transcriptome over several beads, we increase the probability of recovering transcriptomes for all cells. Our platform’s ability to localize combinations of cells, beads, and reagents at defined positions on a nanoliter array affords other powerful capabilities, such as systematic variation of cell, reagent, and drug combinations and tuning of optical and sequencing parameters to achieve optimal data. The open nature of the array also makes it amenable to additional measurement modalities, such as atomic force microscopy, mass spectrometry, and chemical assays, all of which can be linked with optical and scRNA-seq data using the approach we have presented (30,31). The ability to link sequencing readouts with other measurements for thousands of single cells will facilitate further investigations into the molecular underpinnings of cell function (32).

## Materials and Methods

### Microfluidic device fabrication

PDM chips are fabricated by poly(dimethylsiloxane) (PDMS) molding over a SU-8 master. Briefly, a three-layer SU-8 negative master is patterned to form 20, 80, and 150 um tall features using previously described multi-layer SU-8 photolithography techniques (22). Following casting of PDMS over the SU-8 and curing at 65 degrees for 2 hours, inlet holes are punched into devices using a 0.75 mm biopsy core. Devices are then plasma bonded to 25 mm x 75 mm glass slides. 1 cm of PE/5 tubing (Scientific Commodities) is inserted into the nozzle channel and sealed with a 1-minute instant mix epoxy (Norland). Channels are then treated with AquaPel (AquaPel). Drop-making devices are fabricated as previously described (22). Two devices with a T-junction cross-section of 80 um x 45 um and 80 um x 80 um are used.

### Nanoplate fabrication

A negative of the electrode pattern is fabricated on a 50 mm × 75 mm glass slide by positive resist photolithography. A 2 um thick layer of MA-P 1215 (Micro Resist Technology) is spin-coated onto the slide and baked for 1 minute on a 95 degree hotplate. The slide is then exposed to collimated 190 mW UV light (Thorlabs) for 3.5 minutes. The slide is developed in MF-24A developer (Dow Chemical) for 1 minute. Patterned slides then have a 200 A thick layer of chromium deposited on them (LGA Thin Films). The removal of the photoresist with acetone yields the electrode pattern. Nanowells are fabricated on electrode slide by first masking off the regions of electrode contact and spin-coating a 15 um thick layer of uncured PDMS. PDMS is then cured for 3 minutes on a 95 degree hotplate. Following plasma treatment of the slide, a 40 um thick layer of SU-8 is spin coated onto the slide and allowed to soft-bake on a 95 degrees for 10 minutes. The slide is exposed to UV light under a photomask for 90 seconds, followed by 5 minutes of post-exposure baking at 95 degrees. The slide is then immersed in PGMEA developer (Sigma) for 5 minutes, rinsed with PGMEA and isopropanol, then dried on the hot plate for 2 minutes. Slides are then plasma treated and placed in a petri dish adjacent to reservoirs of trichloro(1H,1H,2H,2H-perfluorooctyl)silane (Sigma) for 2 hours under vacuum at room temperature.

### Nanowell coordinate indexing

Nanowells are barcoded using the sciFLEXARRAYER S3 (Scienion AG). A 96 well ‘source-plate’ containing up to 44 coordinate oligos (**Table S1, Table S2**) diluted to 1 nM in DI water is prepared. 2 nL of each barcode oligo solution is dispensed to nanowells according to a pre-programmed print routine to label each nanowell with a unique but known combination of three oligos (**Fig. S1**). Nanowells are split into 14-by-14 subarrays, of which each subarray had 14 unique x and 14 y coordinate oligos. Subarrays are tiled together, with each subarray having a unique z coordinate oligo, until the array reached the desired size. Following printing, slides are placed in a petri dish and sealed with parafilm and stored at −20 degrees until ready to use.

### PDM operation and optical configuration

A multimode excitation fiber with a core diameter of 105 um and a NA of 0.22 (Thorlabs) is inserted into a guide channel in the PDM device. Similarly, an emission detection fiber with core diameter of 200 um and NA of 0.39 (Thorlabs) is inserted into a second guide channel in the PDM device. Four 50 mW continuous wave lasers with wavelengths of 405, 473, 532, and 640 nm are combined and coupled to the excitation fiber. Emitted light is columnated and ported into a quad-bandpass filter, then passed through a series of dichroic mirrors. Bandpass filters of 448, 510, 571, and 697 nm past each dichroic mirror enable wavelength-specific detection of emitted light by PMTs. Electrode channels and a ‘Faraday moat’ are filled with a 5M NaCl solution. A positive electrode is connected to a function generator and a high voltage amplifier while a second electrode is grounded. Fluidic inputs into the PDM device are driven by syringe pumps (New Era). Bias and spacer oil containing 0.2% w/w IK in HFE-7500 are flowed through the device at a flow rate of 2000 uL/h. A waste channel is driven with a negative flow rate of −3000 uL/h. Monodisperse droplet emulsions are reinjected into the device at a flow rate of 100 +/-50 uL/h. Real-time optical signal acquisition through a field programmable gate array (National Instruments) is displayed on a LabView software. Optical signal is processed in real-time and displayed on a fluorescence dot plot, in which drop types of interest can be assigned by specifying gates. Droplets are subsequently sorted by passing a high frequency pulse through a high voltage amplifier (Trek 690E-6). Typical droplet sorting parameters range from 10-20 kHz, 50-100 cycles, and 0.5-1.0 kV. Copper tape with a conductive adhesive (Ted Pella) is affixed to two electrode contact pads on the nanoplate. One pad is connected to ground, while the other one is connected to a function generator and a high voltage amplifier, providing power at 200 - 600V at 20 - 30 kHz. Slides are immersed in a bath of 2% w/w IK in FC-70 (3M) during printing operation.

### Cell culture

HEK and 3T3 cells (ATCC) are cultured in 75 square-cm flasks in the presence of Dulbecco’s Modified Eagle Medium (DMEM) supplemented with 10% fetal bovine serum (FBS) and 1x Penicillin-Streptomycin at 37 deg and 5% CO2. Cells are treated with 0.25% Trypsin-EDTA and ished with media to generate cell supsensions. The viability and cell concentration are counted by a TC20 automated cell counter (BioRad). Cell suspensions are diluted to 1 million/mL in media. Suspensions are pelleted at 400g for 3 minutes and resuspended in 1 mL DPBS. The HEK suspension is treated with 1 ug/mL of Calcein Green (Thermo-Fisher) while the 3T3 suspension is treated with 2 ug/mL of Calcein Red (Thermo-Fisher) for 15 minutes at 37 degrees, followed by the addition of 4 mL media. Suspensions are pelleted and resuspended in media. Cells are mixed together in a 1:1 ratio and diluted in DPBS to form a final concentration of 250k/mL, which contained also 10 uM Cascade Blue-Dextran (Thermo-Fisher) and 0.5 v/v% FBS are added. Jurkat cells (ATCC) are cultured in RPMI-1640 medium supplemented with 10% FBS and 1x Penicillin-Streptomycin at 37 deg and 5% CO2. 1 million cells are extracted and pelleted at 400g for 3 minutes and diluted to a in 500 uL DPBS, to which 1 uL of 5 mM DRAQ5 (Thermo-Fisher) is added. Cells are incubated at 37 degrees for 5 minutes, to which 500 uL of DPBS is added which also contained 10 uM Cascade Blue-Dextran and 0.5 v/v% FBS.

### Cell and bead encapsulation within monodisperse droplet emulsions

Barcoded mRNA capture beads are purchased through ChemGenes (MACOSKO-2011-10) and have a structure previously reported (7). Beads arrived as a dry resin and are resuspended, ished, and filtered as previously described. For each experiment, 100,000 beads are extracted from the suspension and pelleted by placing on a tabletop centrifuge for 10 seconds. The supernatant is removed and replaced with 40 uL of 10 mM 4-MU (Sigma) in methanol diluted in 960 uL DPBS. The pellet is resuspended and allowed to stain for 1 minute at room temperature. Beads are then pelleted, ished with DPBS once, then resuspended in a solution of 10 uM FITC in DPBS to which is added 500 uL of the Drop-Seq lysis buffer. Beads are then placed into a 3 mL syringe with a magnetic stir bar (V&P Scientific) and encapsulated in 2% w/v Ionic Krytox surfactant in HFE 7500 (3M) on an 80 x 80 um drop-making device. Flow rates used are 4000 uL/hr for the bead suspension and 12000 uL/hr for the oil. Cell suspension is placed in a 3 mL syringe with a magnetic stir bar and encapsulated in 2% w/v PEGylated surfactant in HFE 7500 on a 80 x 45 um drop-making device. Flow rates used are 1500 uL/hr for the cell suspension and 4000 uL/hr for the oil.

### PDM operation for performing linked fluorescence and scRNA-seq analysis in nanoplates

The bead-containing droplets are passed through the PDM device at input rates of 80 - 120 Hz. Bead-containing droplets are programmably dispensed to each microwell at a maximum printing rate of 3 Hz between nanowells and 10 Hz if printing multiple beads to the same well. Following printing of beads to nanowells, cell-containing droplets are passed through the PDM device at input rates of 80 - 160 Hz. Cells are dispensed to nanowells at a printing rate of 2-5 Hz. The fluorescence of every cell printed is recorded into a text file along with its nanowell location. Following printing of cells and beads, the nanowell slide is disconnected from its power source, causing droplets to float to the surface, where they are transferred by a P-1000 pipette into a 50 mL conical on ice.

### Sequencing library preparation

The collected emulsions are processed similarly to the Drop-Seq workflow [(7). In brief, the emulsion is broken, beads are collected and reverse transcribed with MMLV reverse transcriptase (Maxima RT, Thermo Fisher), unused primers are degraded with Exonuclease I (New England Biolabs), beads are washed and PCR amplified. The following modifications are incorporated to account for the low number of beads collected. During the emulsion breakage step, a 0.01% v/v solution of Sarkosyl in 6x SSC is used. During the steps leading up to reverse transcription, a 0.01% v/v Tween-20 solution in 6X SSC is used. Following PCR, the cDNA library is split into two fractions following sequential AmPure bead purification at 0.6x and 2.0x volume ratios as performed the Cite-Seq workflow. 600 pg of cDNA in the fraction containing mRNAs is processed using the Nextera XT kit to form a sequencing library. 500 pg of cDNA in the fraction containing amplified well indexes underwent a second round of PCR to add sequencing adapters. Libraries are pooled and sequenced on an Illumina machine.

### NGS sequencing and data analysis

Libraries underwent paired-end sequencing using the custom Drop-Seq primer with a read length of 25 bp for read 1 and 75 bp for read 2. For the mRNA library, reads are processed using the Drop-Seq bioinformatic pipeline. For the well index library, reads are partially processed using the Drop-Seq bioinformatic pipeline, yielding a read-quality filtered and trimmed .sam file with annotations corresponding to UMI and bead barcode positions. Custom Python code is then used to create a well-index expression matrix, with individual beads as the columns and the set of all possible well indexes as the rows, and UMI counts for each possibility of well index and bead populating the cells of the matrix. UMI counts for each bead are scaled based on the number of total UMIs on the bead. Next, the off-target noise of each well index is estimated based on the average expression across all beads and subtracted from scaled UMI counts. The top x, y, and z well index captured on each bead is then extracted. Beads which the top well index is not at least 5 times as abundant as the next most abundant well index for any of the sets of x, y, and z well indexes are removed. The remaining beads are assigned to a nanowell position by matching the most abundant x, y, and z indexes on the bead to a lookup table of the expected x, y, and z positions at each nanowell position. Following position assignment, the bead barcodes of all beads matched at each nanowell position are collected. The columns on the gene expression matrix of all beads matched at the same nanowell position are merged, yielding a revised matrix where the columns represented nanowell positions instead of individual beads. The gene expression matrix is then annotated by recorded cell fluorescence values obtained during printing. For the cell cycle experiment, only those cells which expressed at least 300 genes and could be confidently assigned a fluorescence value are processed using the Seurat package in R. Cells are assigned a G2/M and S phase score using Seurat and a list of previous published cell-cycle associated genes (33) which is then used to assign a cell cycle state. Principal component analysis is performed using only the cell-cycle associated genes and UMAP analysis is then performed on the top 10 principal components.

## Declarations

### Availability of data and materials

Raw sequencing reads as well as nanowell indexed flow cytometry data are available at the Gene Expression Omnibus under accession GSE136871. Python scripts for processing coordinate oligo reads are available in the supplementary material.

### Authors’ contributions

JQZ, CAS, and ARA conceived of the method. JQZ, CAS, LL, and KCC performed experiments. JQZ performed bioinformatic analysis. JQZ, ZJG, and ARA wrote the manuscript. All authors read and approved the manuscript.

## Acknowledgements

We thank the Eric Chow and the Center for Advanced Technology (CAT) at UCSF for providing access to Illumina sequencers and the S3 SciFLEXARRAYER and for assistance with sequencing the Jurkat cell experiment. We thank Christopher McGinnis for helpful discussion regarding bioinformatic data analysis and Russell Cole for support with microfluidic hardware. We thank Sarah Pyle for artistic editing of the figures.

## Supplemental Files

Additional File 1: Python scripts for processing data.

Additional File 2: Supplementary Figures and Tables.

Additional File 3: Movie S1, liftoff of drops following removal of power from electrode array.

Additional File 4: Movie S2, printing of four beads into nanowells, slowed 2x.

