## Additional File 2: Supplemental Figure and Tables for "Linked optical and gene expression profiling of single cells at high throughput"

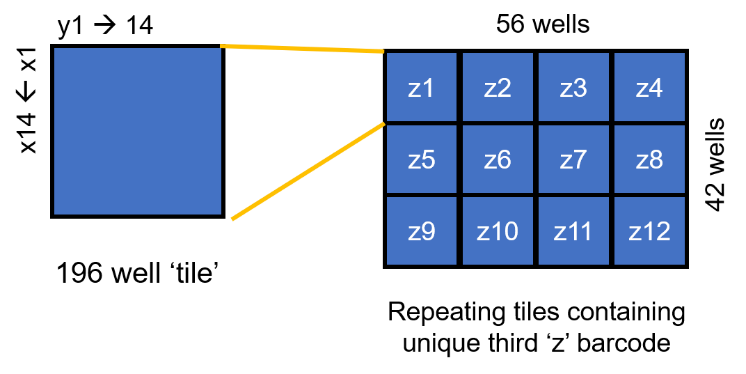

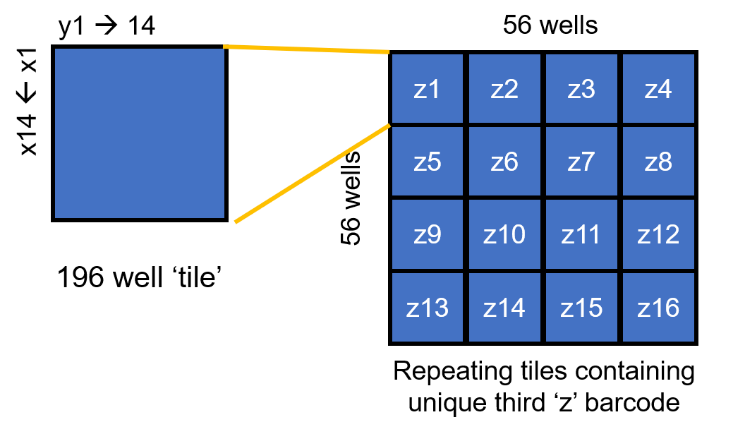

**B**

**A**

**Figure S1**. Well index coordinating scheme. Each well has a unique but known combination of three indexes. A 14 by 14 tile is has each column and row barcoded with a unique index. This tile is repeated a total of (A) twelve or (B) sixteen times with each tile having a unique third index.

| **Component** | **Sequence** |
| --- | --- |
| TSO_PCR Primer | AAGCAGTGGTATCAACGCAGAGTGAAT |
| Illumina Read2 Primer | GTCTCGTGGGCTCGGAGATGTGTATAAGAGACAG |
| Barcode | NNNNNNNN |
| PolyA (21) tail | AAAAAAAAAAAAAAAAAAAAA |

**Table S1**: Components of coordinate oligo, in 5’ to 3’ order.

| **Coordinate Name** | **8bp barcode** |
| --- | --- |
| y1 | CGCCGCAA |
| y2 | ACCAGCCG |
| y3 | TACTGCGA |
| y4 | GTCGGCTG |
| y5 | CCCAGGAC |
| y6 | AACTGGCT |
| y7 | TTCGGGGC |
| y8 | GGCCGGTT |
| y9 | CACTGTAG |
| y10 | ATCGGTCA |
| y11 | TGCCGTGG |
| y12 | GCCAGTTA |
| y13 | GGCATTGT |
| y14 | CCCTTTTC |
| x1 | TGCATACG |
| x2 | GCCTTAGA |
| x3 | CACGTATG |
| x4 | AGCATCAC |
| x5 | TCCTTCCT |
| x6 | GACGTCGC |
| x7 | CTCCTCTT |
| x8 | ACCTTGAG |
| x9 | TACGTGCA |
| x10 | GTCCTGGG |
| x11 | CGCATGTA |
| x12 | TTCCTTCC |
| x13 | TCGTAGTG |
| x14 | GTGCATAC |
| z1 | GGGAAAAG |
| z2 | CCGTAACA |
| z3 | AAGGAAGG |
| z4 | GCGTACAT |
| z5 | CAGGACCC |
| z6 | ATGCACGT |
| z7 | TGGAACTC |
| z8 | GAGGAGAA |
| z9 | CGGAATCT |
| z10 | ACGTATGC |
| z11 | CGGTCAAT |
| z12 | ACGGCACC |
| z13 | TAGCCAGT |
| z14 | GTGACATC |
| z15 | CCGGCCAA |
| z16 | AAGCCCCG |

**Table S2:** Lookup table of the barcode sequences of the coordinate oligos used in this study.
