## Additional File 1: Supplemental Code for "Linked optical and gene expression profiling of single cells at high throughput": array_lut.py

```
import pandas

# converts the excel sheet with the z-coordinates on the grid into a numpy 2d array
coords_LUT = pandas.read_excel('zcoords.xlsx', sheet_name='Sheet4', header=None).values
rows = coords_LUT.shape[0]
cols = coords_LUT.shape[1]

# adds the proper x and y barcodes for each position
for i in range(rows):
    for ii in range(cols):
        coords_LUT[i, ii] = str(['x' + str(ii % 14 + 1), 'y' + str(i % 14 + 1), coords_LUT[i, ii]])

# create lookup table for x,y,z barcode --> 8 BP sequence
df = pandas.read_excel('zcoords.xlsx', sheet_name='probes to plate', header=None).values
xyz_LUT = {value[1]: value[0] for value in df}
```
