## Additional File 1: Supplemental Code for "Linked optical and gene expression profiling of single cells at high throughput": bead_barcodes_anal_Jul2019.py

```
from matrix_parse import barcode_positions
from gen_mixed_species_plot import beads
# from extract_file_names_for_cellmatrix import cells_matrix
import numpy as np
import matplotlib.pyplot as plt
import pandas as pd

def array_coord_to_pos_no(coord):
    '''
    :param coord: a tuple contatining (row, column) zero-indexed on a printed droplet array
    :return: the position number of coordinate
    '''
    row = coord[0]
    col = coord[1]
    if not row % 2:
        pos_no = row * 56 + col + 1
    else:
        pos_no = (row + 1) * 56 - col
    return pos_no
def matchScore(string1, string2):
    score = 0
    for char1, char2 in zip(string1, string2):
        if char1 == char2:
            score += 1
    return score

"""
Create plot of number of transcripts per well
"""
correct_species_arr = np.full((56, 56), 0)
missing_keys = 0
inc_key = 0
beads_mapped = 0

trans_per_well = {}
short_frac_keys = [key for key in barcode_positions.keys()]

for key in beads.keys():
    # foundKey = False
    if key not in barcode_positions:
        #small modification to allow for a hamming distance of 1 when considering short and long fraction barcode matches
        #if hamming distance >= 11 (unused)
        # for key2 in short_frac_keys:
        #     if matchScore(key, key2) == 11:
        #         barcode_positions[key] = barcode_positions[key2]
        #         if barcode_positions[key2] in trans_per_well:
        #             trans_per_well[barcode_positions[key2]] = (trans_per_well[barcode_positions[key2]][0] + beads[key][0],
        #                                                     trans_per_well[barcode_positions[key2]][1] + beads[key][1])
        #         else:
        #             trans_per_well[barcode_positions[key2]] = beads[key]
        #         foundKey = True
        #         beads_mapped += 1
        #         break
        missing_keys += 1
    else:
        beads_mapped += 1
        # concatenate info if several beads are mapped to the same spot
        if barcode_positions[key] in trans_per_well:
            trans_per_well[barcode_positions[key]] = (trans_per_well[barcode_positions[key]][0] + beads[key][0],
                                                    trans_per_well[barcode_positions[key]][1] + beads[key][1])
        else:
            trans_per_well[barcode_positions[key]] = beads[key]

keys_to_remove = []
for key in trans_per_well.keys():
    correct_species_arr[key[0]][key[1]] = sum(trans_per_well[key])

#     # optional for transcript-based filtering out of wells
#     if sum(trans_per_well[key]) < 800:
#         keys_to_remove.append(key)
#
# for key in keys_to_remove:
#     del(trans_per_well[key])


plt.figure(1)
num_wells_mapped = len([key for key in trans_per_well.keys()])
print(str(beads_mapped) + ' beads mapped and ' + str(num_wells_mapped) + ' wells mapped')
heatmap = plt.imshow(correct_species_arr, cmap='hot')
bar = plt.colorbar()
# bar.set_ticks([-0.5, 0, .5, 1])
# bar.set_ticklabels(['empty well', 'human', 'mixed', 'mouse'])
# bar.set_label('Composition of transcripts within microwell')
plt.xlabel('Array position (column)')
plt.ylabel('Array position (row)')


"""
Correlate wells with recovered transcripts to imaging ground truth 
"""

trans_recovered = []  # keeps track of transcripts per well if there was a cell originally in the well
missing_image_but_RNA = []  # keeps track of which wells have mRNA captured but no imaged cell
no_bead_mapped = 0
species_mapping_correctness = np.full((56, 56), -0.5)
inc_key = 0
for i in range(0, species_mapping_correctness.shape[0]):
    for j in range(0, species_mapping_correctness.shape[1]):
        if (i, j) in trans_per_well.keys():
            if sum(trans_per_well[i, j]) < 1:
               continue
            human_species_purity = trans_per_well[(i, j)][0] / sum(trans_per_well[i, j])
            species_mapping_correctness[i, j] = human_species_purity
            trans_recovered.append(sum(trans_per_well[i, j]))
            if not (i + j) % 2:
                if not human_species_purity < 0.1:  # mouse cell
                    inc_key += 1
            else:  # mod2 means human cell
                if not human_species_purity > 0.9:  # human cell
                    inc_key += 1

# for key in trans_per_well.keys():
#     if cells_matrix[key[0], key[1]] > 0 and sum(trans_per_well[key]) > 0:
#         species_mapping_correctness[key[0], key[1]] = 1
#     elif sum(trans_per_well[key]) > 0 and cells_matrix[key[0], key[1]] == 0:
#         species_mapping_correctness[key[0], key[1]] = -1

num_wells_mapped = len([v for v in species_mapping_correctness.flatten() if v != -0.5])

print('incorrect wells: ' + str(inc_key) + ' of ' + str(num_wells_mapped) + ' total wells.')

plt.figure(2)
heatmap = plt.imshow(species_mapping_correctness, cmap='hot')
bar = plt.colorbar()
bar.set_ticks([-0.5, 0, .5, 1])
bar.set_label('Human species purity (black = empty well)')
plt.title('matching of image to RNA-seq by microwell (green = correct)')
plt.xlabel('Array position (column)')
plt.ylabel('Array position (row)')


plt.figure(3)
plt.hist(trans_recovered, bins=100)
plt.ylabel('counts')
plt.xlabel('number of transcripts recovered from well')


plt.show()

'''
Output of mapped wells to excel files, for further analyses


# for key in barcode_positions.keys():
#     barcode_positions[key] = array_coord_to_pos_no(barcode_positions[key])
# df = pd.DataFrame(barcode_positions, index=[0])
# writer = pd.ExcelWriter('cell_barcode_positionsAug2019.xlsx')
# df.to_excel(writer)
# writer.save()

keys_to_remove = []
for key in barcode_positions.keys():
    if key not in beads.keys():
        keys_to_remove.append(key)

for key in keys_to_remove:
    del(barcode_positions[key])

# output which wells were mapped to an excel file
mapped_array_out = {}

for key in barcode_positions.keys():
    if array_coord_to_pos_no(barcode_positions[key]) in mapped_array_out:
        mapped_array_out[array_coord_to_pos_no(barcode_positions[key])].append(key)
    else:
        mapped_array_out[array_coord_to_pos_no(barcode_positions[key])] = [key]

# remove entries from the dictionary that correspond to wells that have RNA linked but no imaged cell
for position in missing_image_but_RNA:
    del(mapped_array_out[position])

df2 = pd.DataFrame.from_dict(data = mapped_array_out, orient='index')
df2.to_excel('mapped_bead_barcodes_per_well.xlsx')
'''
```
