## Additional File 1: Supplemental Code for "Linked optical and gene expression profiling of single cells at high throughput": convert_customreads_to_matrix.py

```
import numpy as np
import matplotlib.pyplot as plt
import csv
import sys
import pandas as pd
from math import log

csv.field_size_limit(20000000)


# read noUMIs file first from short fraction
with open('dict_corrected_noUMIs', mode='r') as file:
    reader = csv.reader(file)
    barcode_groups = {}

    progress = 0 # iteration tracker
    for row in reader:

        # progress counter
        if not progress % 1000:
            print(str(progress) + ' barcode groups read')
        progress += 1

        bead_barcode, reads = row  # each ROW has two strings, a key-value pair
        reads = eval(reads)  # required to unpack VALUE, which is a string representation of a dictionary

        barcode_groups[bead_barcode] = reads

# read in well indexes, and make a dict of the format {8BP_barcode: coordinate, ...}
data = pd.read_csv(sys.argv[2], header=None)
well_index_LUT = {}
for item, row in data.iterrows():
    well_index_LUT[row[0]] = row[1]


# make dict of the format: {barcode: [x1counts, x2counts, ..., z16 counts]}
well_index_names = ['x1', 'x2', 'x3', 'x4', 'x5', 'x6', 'x7', 'x8', 'x9', 'x10', 'x11', 'x12', 'x13', 'x14',
                  'y1', 'y2', 'y3', 'y4', 'y5', 'y6', 'y7', 'y8', 'y9', 'y10', 'y11', 'y12', 'y13', 'y14',
                  'z1', 'z2', 'z3', 'z4', 'z5', 'z6', 'z7', 'z8', 'z9', 'z10', 'z11', 'z12', 'z13', 'z14',
                  'z15', 'z16']

to_df = {}
counts = []   # for plotting distribution of nUMI counts for all barcode groups
progress = 0


for barcode in barcode_groups.keys():
    progress += 1
    nUMIs_dict = {} # contains a dict: {coordinate: nUMIs, ...}
    temp = barcode_groups[barcode]   # temp just contains a dictionary of the form {8bpbarcode: nUMIs, ...}
    for key in temp.keys():
        nUMIs_dict[well_index_LUT[key]] = temp[key]

    lst = []  # contains a list of UMI counts for each well_index: [nUMI x1, nUMI x2, ...]
    for well_index in well_index_names:
        if well_index in nUMIs_dict:
            lst.append(nUMIs_dict[well_index])
        else:
            lst.append(0)

    if not progress % 1000:
        print(str(progress) + ' barcode groups read')
    if sum(lst) > 90:
        to_df[barcode] = lst
    # lst.sort(key=lambda x: x[0])
    if sum(lst) < 10:
        continue
    counts.append(log(sum(lst)))


"""
From plotting histogram of UMI counts, a reasonable cutoff for what is considered a 'real' bead is about 90 UMIs
"""
# plt.hist(counts, bins='fd')
# plt.show()

df = pd.DataFrame.from_dict(data=to_df, orient='index', columns=well_index_names)
print(df)
df['nUMIs'] = df.apply(sum, axis=1)   # uses df.sum to append a column of nUMIs

df.to_excel("out.xlsx")
```
