## Additional File 1: Supplemental Code for "Linked optical and gene expression profiling of single cells at high throughput": correct_bead_synthesis_errors.py

```
'''
Corrects for bead synthesis errors:

Looks at position 8 of the UMI for given bead barcode groups. If the frequency of T at position 8 is over 80%:
    - cut out base 8 of the UMI
    - move base 12 of the bead barcode over to base 1 of the UMI
    - move base 1 - 7 of the UMI to base 2 - 8
    - replace base 12 of the bead barcode with 'N'

Outputs a corrected form of the dictionary:
    {bead_code: {well_coord1: UMI's}...}
'''

import sys
import csv
import matplotlib.pyplot as plt
from collections import Counter

csv.field_size_limit(80000000)

### first generate {bead_code: {well_coord1: UMI's}...} dictionary

with open('dict_hamming1', mode='r') as file:
    reader = csv.reader(file)
    barcode_groups = {}
    for row in reader:
        key, value = row  # each ROW has two strings, a key-value pair
        value = eval(value)  # required to unpack VALUE, which is a string representation of a dictionary
        barcode_groups[key] = value

corrected_dict = {}

def count_character_frequencies(lst_of_strings):
    '''
    Takes a lst_of_strings of strings of the same length, and returns a list where each value is a dictionary containing
    the character frequencies at that character position
    '''
    output = []
    lst_of_chars = list(zip(*lst_of_strings)) # creates a list where each entry contains the nth character for every
        # single string
    total_strings = len(lst_of_strings)
    for chara_lst in lst_of_chars:
        c = Counter()
        for chara in chara_lst:
            c[chara] += 1 / total_strings
        output.append(dict(c))
    return output

### iterate over all bead barcode groups, IF it has at least 15 UMI's
keys_to_be_deleted = []
for key in barcode_groups.keys():
    UMIs = []
    for well_coord in barcode_groups[key]:
        UMIs.extend(barcode_groups[key][well_coord])  # should now contain all UMI's for a bead barcode
    if len(UMIs) < 15:  # IF it has at least 15 UMI's
        corrected_dict[key] = barcode_groups[key]
        continue
    base_freqs = count_character_frequencies(UMIs)

    # look for a frequency of T at position 8 of the UMI of at least 0.8, and correct UMIs and bead barcode
    if 'T' in base_freqs[7] and base_freqs[7]['T'] > 0.8:
        print('w3w')
        new_key = key[:11] + 'N'
        # change all UMI's in dictionary
        dict_of_UMIs = barcode_groups[key]
        for well_coord in dict_of_UMIs.keys():
            new_UMI_lst = []
            for UMI in dict_of_UMIs[well_coord]:
                new_UMI_lst.append(key[11] + UMI[:7])
            dict_of_UMIs[well_coord] = new_UMI_lst

        # dict_of_UMIs now contains {well_coord: [corrected UMIs];...}

        # create a new entry in corrected_dict corresponding to the corrected key and UMIs
        if new_key in corrected_dict:  # if the key already exists, see if the well_coords exist
            for well_coord in dict_of_UMIs:
                if well_coord in corrected_dict[new_key]:
                    corrected_dict[new_key][well_coord].extend(dict_of_UMIs[well_coord])
                else:
                    corrected_dict[new_key][well_coord] = dict_of_UMIs[well_coord]
        else:
            corrected_dict[new_key] = dict_of_UMIs
    else:
        corrected_dict[key] = barcode_groups[key]

# with open('Short Fraction/dict_short_corrected_hamming1', 'w', newline='') as csv_file:
#     writer = csv.writer(csv_file)
#     for key, value in corrected_dict.items():
#         writer.writerow([key, value])

# writes a file of {bead_barcode: {well_coord: # reads};...}

'''
counts all unique UMI's in each well barcode group of each bead barcode
'''

UMI_dist = [] # variable that will be used to generate a frequency plot of all UMI counts
for key in corrected_dict.keys():
    for key2 in corrected_dict[key].keys():
        corrected_dict[key][key2] = len(set(corrected_dict[key][key2]))
        UMI_dist.append(corrected_dict[key][key2])

with open('dict_corrected_noUMIs', 'w', newline='') as csv_file:
    writer = csv.writer(csv_file)
    for key, value in corrected_dict.items():
        writer.writerow([key, value])
```
