## Additional File 1: Supplemental Code for "Linked optical and gene expression profiling of single cells at high throughput": extract_bead_barcode_groups.py

```
import csv
import random
import sys

'''
Custom Python Code for "Linked optical and gene expression profiling of single cells at high throughput"

The well index fraction reads must be processed through to the 'SamToFastq' step of the Drop-seq alignment pipeline
'''

def matchScore(string1, string2):
    score = 0
    for char1, char2 in zip(string1, string2):
        if char1 == char2:
            score += 1
    return score

def if_hamming_match(string, string_lst, score):
    '''
    Since all the well barcodes are at least a hamming distance of 3 apart, reads that have a hamming distance of 1
    with respect to a known barcode can be corrected for
    :param string: input string
    :param string_lst: list of well barcodes
    :param score: minimum matchscore of string to any string in string_lst
    :return: a matched string with a match score of at least SCORE, otherwise None
    '''
    for str in string_lst:
        if matchScore(string, str) >= score:
            return str
    return None

# set of bead barcodes
well_barcodes = {'CGCCGCAA','ACCAGCCG','TACTGCGA','GTCGGCTG','CCCAGGAC','AACTGGCT','TTCGGGGC','GGCCGGTT','CACTGTAG',
                 'ATCGGTCA','TGCCGTGG','GCCAGTTA','TGCATACG','GCCTTAGA','CACGTATG','AGCATCAC','TCCTTCCT','GACGTCGC',
                 'CTCCTCTT','ACCTTGAG','TACGTGCA','GTCCTGGG','CGCATGTA','TTCCTTCC','GGCATTGT','CCCTTTTC','GGGAAAAG',
                 'CCGTAACA','AAGGAAGG','GCGTACAT','CAGGACCC','ATGCACGT','TGGAACTC','GAGGAGAA','TCGTAGTG','GTGCATAC',
                 'CGGAATCT','ACGTATGC','CGGTCAAT','ACGGCACC','TAGCCAGT','GTGACATC','CCGGCCAA','AAGCCCCG'}
barcode_groups = {}

progress = 0 # loop iteration tracker
# ZIP() allows you to access two iterables in parallel at once
file = open('YOUR .SAM FILE', 'r')
i = 0
for line in file:
    i += 1
    # # only sample 1/10 of reads
    # if random.random() > 0.1:
    #     continue
    # first 2 lines of the .sam file are headers
    if i < 3:
        continue

    # .sam file has gone through DropSeq pipeline up to TrimSmartAdapater
    # .sam file has some additional annotations at the end of the line corresponding to the bead barcode and UMI
    record_split_by_whitespace = line.split()
    well_barcode = record_split_by_whitespace[9][0:8]
    bead_barcode = record_split_by_whitespace[11][-12:]
    UMI = record_split_by_whitespace[13][-8:]

    # skip next steps if not a valid well barcode
    # this will only consider ~80% of all reads; with Hamming-correction, we can get 15% of lost reads back
    well_barcode = if_hamming_match(well_barcode, well_barcodes, 7)
    if not well_barcode:
        continue

    # progress tracker; updates every 10000 sequences read
    if not progress % 10000:
        print(str(progress) + ' sequences read')
    progress += 1

    # make dictionary of the form {bead barcode: {coord1: reads; coord2: reads}; ...}
    # if bead_barcode in barcode_groups:
    #     if well_barcode in barcode_groups[bead_barcode]:
    #         barcode_groups[bead_barcode][well_barcode] += 1
    #     else:
    #         barcode_groups[bead_barcode][well_barcode] = 1
    # else:
    #     barcode_groups[bead_barcode] = {well_barcode: 1}

    # # # # records individual UMI's
    if bead_barcode in barcode_groups:
        if well_barcode in barcode_groups[bead_barcode]:
            barcode_groups[bead_barcode][well_barcode].append(UMI)
        else:
            barcode_groups[bead_barcode][well_barcode] = [UMI]
    else:
        barcode_groups[bead_barcode] = {well_barcode: [UMI]}

# # counts all unique UMI's in each well barcode group of each bead barcode
# UMI_dist = [] # variable that will be used to generate a frequency plot of all UMI counts
# for key in barcode_groups.keys():
#     for key2 in barcode_groups[key].keys():
#         barcode_groups[key][key2] = len(set(barcode_groups[key][key2]))
#         UMI_dist.append(barcode_groups[key][key2])
#

# writes the output to a csv file where each row is a bead barcode group
with open('dict_hamming1', 'w', newline='') as csv_file:
    writer = csv.writer(csv_file)
    for key, value in barcode_groups.items():
        writer.writerow([key, value])
```
