## Additional File 1: Supplemental Code for "Linked optical and gene expression profiling of single cells at high throughput": gen_mixed_species_plot.py

```
import numpy as np
import matplotlib.pyplot as plt
import csv

'''
For Jurkats Oct 2018
'''

f1 = open('out_cell_readcounts.txt', 'r')

beads = {}  # dictionary of the form {cell_barcode: [# human, # mouse transcripts]...}
for file in [f1]:
    lineno = 0
    for line in file:  # contains info about beads that have human transcripts
        lineno += 1
        if lineno > 2000: # these lines don't contain useful information
            continue
        else:
            num_reads, cell_barcode = line.split()
            num_transcripts = num_reads
            beads[cell_barcode] = [int(num_transcripts), 0]

# writes the output to a csv file where each row is a bead barcode group
# with open('dict_mixed_species.csv', 'w') as csv_file:
#     writer = csv.writer(csv_file)
#     for key, value in beads.items():
#         writer.writerow([key, value])


# filter dictionary by removing all bead barcode groups with less than N transcripts
# OR less than 80% species-purity
keys_to_remove = []
for key in beads.keys():
    if sum(beads[key]) < 200:
        keys_to_remove.append(key)
    # elif beads[key][0] > beads[key][1]:
    #     if beads[key][1]/beads[key][0] > 0.2:
    #         keys_to_remove.append(key)
    # elif beads[key][1] > beads[key][0]:
    #     if beads[key][0]/beads[key][1] > 0.2:
    #         keys_to_remove.append(key)

keys_to_remove = set(keys_to_remove)
for key in keys_to_remove:
    del(beads[key])

print(len([key for key in beads.keys()]))

'''
Well index matching

well_barcodes = {'TACGTGCA', 'AGGGGAAA'}


# read in well_coordinate files:
# well_coord_groups is a DICT that has format {bead_barcode: {indexA: #UMI's}, bead_barcode{indexB: #UMI's}, ...}
well_coord_groups = {}
for fname in ['Short Fraction/dict_short_corrected_noUMIs']:
    with open(fname, mode='r') as file:
        reader = csv.reader(file)
        for row in reader:
            key, value = row  # each ROW has two strings, a key-value pair
            value = eval(value)  # required to unpack VALUE, which is a string representation of a dictionary
            well_coord_groups[key] = value

def matchScore(string1, string2):
    score = 0
    for char1, char2 in zip(string1, string2):
        if char1 == char2:
            score += 1
    return score

def keywithmaxval(d):
     """ a) create a list of the dict's keys and values;
         b) return the key with the max value"""
     v=list(d.values())
     k=list(d.keys())
     return k[v.index(max(v))]


# score each key in BEADS against all keys in well_coord_groups UNTIL you get a key with a score = 12
for key1 in beads.keys():
    for key2 in well_coord_groups.keys():
        if matchScore(key1, key2) == 12:
            majority_well_coord = keywithmaxval(well_coord_groups[key2])
            ###
            # Remove all beads that have a well coord purity of less than 1 - 1/(x + 1)
            ###
            if majority_well_coord == 'TACGTGCA' and 'AGGGGAAA' in well_coord_groups[key2]:
                if well_coord_groups[key2]['TACGTGCA']/well_coord_groups[key2]['AGGGGAAA'] < 4:
                    break
            elif majority_well_coord == 'AGGGGAAA' and 'TACGTGCA' in well_coord_groups[key2]:
                if well_coord_groups[key2]['AGGGGAAA']/well_coord_groups[key2]['TACGTGCA'] < 4:
                    break
            beads[key1].append(keywithmaxval(well_coord_groups[key2]))

# filter dictionary by removing all bead barcode groups that didn't map to a well coord
keys_to_remove = []
counter = 0
counter2 = 0
for key in beads.keys():
    if len(beads[key]) == 2:
        counter2 += 1
        keys_to_remove.append(key)

keys_to_remove = set(keys_to_remove)
print(len(keys_to_remove))
for key in keys_to_remove:
    del(beads[key])
'''

'''
Plotting with well index annotation


# create list of [[# human, # mouse], [# human, # mouse]...] transcripts for each well coord
well_barcodes = ['TACGTGCA', 'AGGGGAAA']
transcript_counts_red = [beads[key][:2] for key in beads.keys() if beads[key][2] == well_barcodes[0]]
transcript_counts_blue = [beads[key][:2] for key in beads.keys() if beads[key][2] == well_barcodes[1]]

# convert list to two sets of ordered lists for scatter plot
human_transcripts_red, mouse_transcripts_red = zip(*transcript_counts_red)
human_transcripts_blue, mouse_transcripts_blue = zip(*transcript_counts_blue)

plt.scatter(human_transcripts_red, mouse_transcripts_red, color='red')
plt.scatter(human_transcripts_blue, mouse_transcripts_blue, color='blue')
axes = plt.gca()
axes.set_xlim([-200, 10000])
axes.set_ylim([-200, 20000])
plt.xlabel('Human transcripts')
plt.ylabel('Mouse transcripts')
lgnd = plt.legend(['Well index = TACGTGCA (n = %i)' % len(transcript_counts_red),
                   'Well index = AGGGGAAA (n = %i)' % len(transcript_counts_blue)])
lgnd.get_frame().set_edgecolor('black')
plt.show()
'''

'''
Plotting (no well index)


# create list of [[# human, # mouse], ...] transcripts for each bead
transcript_counts = [beads[key] for key in beads.keys()]

# replace all zeros with 0.01
transcript_counts2 = []
for x in transcript_counts:
    if x[0] == 0:
        transcript_counts2.append([0.01, x[1]])
    elif x[1] == 0:
        transcript_counts2.append([x[0], 0.01])
    else:
        transcript_counts2.append(x)

human_beads = [x for x in transcript_counts2 if x[1]/x[0] < 0.2 and x[0] > x[1]]
mouse_beads = [x for x in transcript_counts2 if x[0]/x[1] < 0.2 and x[1] > x[0]]
mixed_beads = [x for x in transcript_counts2 if x not in human_beads and x not in mouse_beads]
print(len(human_beads), len(mixed_beads), len(mouse_beads))

#human_beads_human_trans ~ hh; human_beads_mouse_trans ~ hm
hh, hm = zip(*human_beads)
mh, mm = zip(*mouse_beads)
mixh, mixm = zip(*mixed_beads)

# human_transcripts, mouse_transcripts = zip(*transcript_counts)
plt.figure(figsize=(6.5, 5))
plt.scatter(hh, hm, color='red')
plt.scatter(mh, mm, color='blue')
plt.scatter(mixh, mixm, color='purple')
plt.legend(('Human: N = ' + str(len(human_beads)),
            'Mouse: N = ' + str(len(mouse_beads)),
            'Mixed: N = ' + str(len(mixed_beads))))
axes = plt.gca()
axes.set_xlim([-500, 20000])
axes.set_ylim([-500, 15000])
plt.xlabel('Human transcripts')
plt.ylabel('Mouse transcripts')
plt.rc('font', size=20)
plt.tight_layout()
plt.show()

'''
```
