## Additional File 1: Supplemental Code for "Linked optical and gene expression profiling of single cells at high throughput": matrix_parse.py

```
import pandas as pd
import numpy as np
import matplotlib.pyplot as plt
import sys
from array_lut import coords_LUT

# import cite-seq matrix
core_matrix = pd.read_excel('out.xlsx', sheet_name='baselinesub', index_col=0)
print('core matrix loaded')

barcode_positions = {}
num_matched_beads = 0

# core_matrix = core_matrix[:4000]

# iterates over rows in matrix, which are labeled by bead barcode
for barcode, row in core_matrix.iterrows():
    top_indexes = []

    # extract x, y, and z barcode abundances in this format:
    # x_abundances = [('x1', 0), ('x2', 2) ... ]
    x_abundances = [('x1', row['x1']), ('x2', row['x2']), ('x3', row['x3']), ('x4', row['x4']), ('x5', row['x5']),
                    ('x6', row['x6']), ('x7', row['x7']), ('x8', row['x8']), ('x9', row['x9']), ('x10', row['x10']),
                    ('x11', row['x11']), ('x12', row['x12']), ('x13', row['x13']), ('x14', row['x14'])]
    y_abundances = [('y1', row['y1']), ('y2', row['y2']), ('y3', row['y3']), ('y4', row['y4']), ('y5', row['y5']),
                    ('y6', row['y6']), ('y7', row['y7']), ('y8', row['y8']), ('y9', row['y9']), ('y10', row['y10']),
                    ('y11', row['y11']), ('y12', row['y12']), ('y13', row['y13']), ('y14', row['y14'])]
    z_abundances = [('z1', row['z1']), ('z2', row['z2']), ('z3', row['z3']), ('z4', row['z4']), ('z5', row['z5']),
                    ('z6', row['z6']), ('z7', row['z7']), ('z8', row['z8']), ('z9', row['z9']), ('z10', row['z10']),
                    ('z11', row['z11']), ('z12', row['z12']), ('z13', row['z13']), ('z14', row['z14']),
                    ('z15', row['z15']), ('z16', row['z16'])]

    # sort each of these
    for abundance_lst in [x_abundances, y_abundances, z_abundances]:
        abundance_lst = sorted(abundance_lst, key=lambda x: x[1], reverse=True)

    # if top barcode is at least 2.5x more abundant than the next one, record it
    # but barcode needs to be at least 5 umi's
        if abundance_lst[0][1] > 3 and abundance_lst[0][1] > abundance_lst[1][1] * 3:
            top_indexes.append(abundance_lst[0][0])
    if len(top_indexes) == 3:
        barcode_positions[barcode] = top_indexes
        num_matched_beads += 1

print(str(num_matched_beads) + ' beads matched to position')

keys_to_remove = []

# replace each value in barcode_positions with array_position, else delete it
for barcode in barcode_positions.keys():
    if str(barcode_positions[barcode]) in coords_LUT:
        barcode_positions[barcode] = list(zip(*np.where(coords_LUT == str(barcode_positions[barcode]))))[0]
    else:
        keys_to_remove.append(barcode)

for barcode in keys_to_remove:
    del(barcode_positions[barcode])

""""
CREATE PLOT OF MATCHED BEADS TO ARRAY POSITION
"""

# initalize a zeroes array the size of the array
heatmap_arr = np.zeros((56, 56))

matched_beads = 0
# for each matched position on the array, increase the value in HEATMAP_ARR by 1
for key in barcode_positions.keys():
    matched_beads += 1
    heatmap_arr[barcode_positions[key][0], barcode_positions[key][1]] += 1

print(str(matched_beads) + ' beads mapped to an array position')
plt.figure(1)
heatmap = plt.imshow(heatmap_arr, cmap='hot')
bar = plt.colorbar()
bar.set_ticks([0, 1, 2, 3, 4, 5, 6, 7])
bar.set_label('Number of beads mapped to position')
plt.xlabel('Array position (column)')
plt.ylabel('Array position (row)')
# plt.colorbar(heatmap)

# from collections import Counter
# print(Counter(heatmap_arr.flatten()))

print(len([v for v in heatmap_arr.flatten() if v > 0]))
plt.figure(2)
n, bins, patches = plt.hist(heatmap_arr.flatten(), bins='auto')  # [1, 2, 3, 4, 5, 6, 7])
print(n)
plt.xlabel('number of times a bead barcode matches to a particular well')
plt.ylabel('frequency')
plt.show()
```
