## Additional File 1: Supplemental Code for "Linked optical and gene expression profiling of single cells at high throughput": merge_expression_matrix.py

```
import pandas as pd

"""
July2019
"""

'''
Turns a DGE that has cell barcodes as columns to a DGE with well No. as columns
'''


# read in the DGE as a pandas dataframe
dge_matrix = pd.read_csv('cleaned_star_gene_exon_tagged.dge.txt', delimiter='\t', header=0, index_col=0)
#print(dge_matrix)

# read in mapped_wells into dataframe
mapped_barcodes_by_well = pd.read_excel('mapped_bead_barcodes_per_well.xlsx') # .transpose()  # columns are well No.

# for some reason, Pandas has a weird NaN that needs to be replaced
mapped_barcodes_by_well = mapped_barcodes_by_well.replace({pd.np.nan: None})

# initialize an empty DataFrame (gene expression matrix)
dge_combined = pd.DataFrame(index=dge_matrix.index)

mapped_barcodes_by_well = mapped_barcodes_by_well.values.tolist()
for well in mapped_barcodes_by_well:
    well_no = well[0]
    for item in well[1:]:
        if item:
            barcode = item
            if well_no not in dge_combined.columns.values:
                dge_combined[well_no] = dge_matrix[barcode]
            else:
                if barcode in dge_matrix.columns.values:
                    dge_combined[well_no] += dge_matrix[barcode]

    # for barcode in mapped_barcodes_by_well[well_no]:
    #     print(barcode)
    #     if barcode:
    #         if well_no not in dge_combined.columns.values:
    #             dge_combined[well_no] = dge_matrix[barcode]
    #         else:x
    #             if barcode in dge_matrix.columns.values:
    #                 dge_combined[well_no] += dge_matrix[barcode]

dge_combined.to_csv('combined_dge.csv')
print(dge_combined)
```
