## Additional File 1: Supplemental Code for "Linked optical and gene expression profiling of single cells at high throughput": README.txt

Custom Python Code for "Linked optical and gene expression profiling of single cells at high throughput"

The well index fraction reads must be processed through to the 'SamToFastq' step of the Drop-seq alignment pipeline.

Requires Python 3
Requires Drop Seq analysis tools v1.x.x

1. Run extract_bead_barcode_groups.py  with [.sam filename] on (line42)
This will group together all beads based on bead barcode, and will group together all reads associated with the well barcodes with a hamming correction of 1.

2. Run correct_bead_synthesis_errors.py with dict_hamming1 on (line23)
This will perform bead barcode correction as outlined in Drop-Seq: for beads with at least 15 UMIs associated, if the frequency at position 8 of the UMI is at least 80% T, it will shift barcode position #12 to UMI position #1, delete UMI position #8, and replace barcode position #12 with N.

3. Run convert_customreads_to_matrix.py with dict_corrected_noUMIs on (line12)
This will convert the reads into a matrix of bead barcodes and expression of well indexes. 
After step 3, there will be an excel file output called 'out.xlsx'

To this file: normalize each cell by number of total UMIs for that bead, then multiply by 10000
For each cell, subtract the average expression of that well index across all beads, and set the cell's value to 0 if this yields a negative number. 

Store this modified matrix in a sheet called 'baselinesub' 

4. Ensure that both output files from the Drop-Seq pipeline are available.
Run 'bead_barcodes_anal_Jul2019.py'
Line9 of gen_mixed_species_plot.py might need to be adjusted to match the dge.summary.txt file output 
Transcripts thresholds can be set in gen_mixed_species_plot.py on line34

This will output a file that contains a matrix of well positions, and all bead barcodes associated with that well

5. Run merge_expression_matrix.py
Set line13 to be the digital gene expression matrix
Set line17 to be the outfile from Step 4. 

This will output a corrected gene expression matrix that will only contains wells for which there are associated beads. This expression matrix can be analyzed using R, or other DGE analysis tools. 
